# Geometric Study in Binocular Vision of Healthy Human Eye’s Misaligned Optical Components

**DOI:** 10.1101/2024.06.01.595622

**Authors:** Jacek Turski

## Abstract

The healthy human eye’s optical components are misaligned, which has been overlooked in studies of binocular vision. This study presents comprehensive geometric constructions in the binocular system, with the eye model incorporating the fovea that is not located on and the lens that is tilted away from the eye’s optical axis. It includes the vertical misalignment of optical components and the 3D binocular field of fixations with the previously considered horizontal misalignment underlying stereopsis. The consequences and functional roles of vertical misalignment are explained in the following finding: The classic Helmholtz theory, which states that the subjective vertical retinal meridian inclination to the retinal horizontal meridian explains the perceived backward tilt of the vertical horopter, is less relevant when the eye’s optical components are misaligned. Instead, the lens vertical tilt provides the retinal vertical criterion. The typical misalignment of the fovea and lens in the human population confirms the experimentally measured vertical horopter’s inclination.

## 1 Introduction

The optical components of the healthy human eyes are misaligned; the fovea is not located on, and the crystalline lens is tilted away from the eye’s optical axis. This asymmetry has been accounted for in many clinical studies, usually decomposed along temporal-nasal and inferior-superior axes, that is, into horizontal and vertical segments, respectively [Chang et al., 2007, de Castro et al., 2007, Schaeffel and Kaymak, 2010, Aguirre, 2019, Wang et al., 2019].

The consequences of the horizontal misalignment of the eye’s optical components for the quality of vision are well known—the fovea’s anatomical displacement in the retina from the posterior pole contributes to optical aberrations, and the lens tilt partially compensates for these aberrations [Tabernero et al., 2007, Charman and Atchison, 2009, Artal, 2014, Liu and Thibos, 2017].

To introduce another consequence of this horizontal misalignment, I first recall the notion of stereopsis. If for two retinal elements, one in each eye, a localized stimulus is perceived in a single direction irrespective of whether it reaches only one element or the other or both simultaneously, they are considered to be corresponding elements of zero disparity determined by the nonius paradigm [Noorden von and Campos, 2002]. The locus of points in the binocular field, such that each point projects to a pair of corresponding retinal elements, is known as the horopter. The horizontal correspondence between disparate 2D retinal images of our laterally separated two eyes *organizes* visual perception of objects’ form and their location in 3D space, i.e., stereopsis and spatial relations [Wheatstone, 1838, Julesz, 1971].

The measurements revealed the asymmetry of the horizontal distribution of retinal corresponding elements: the corresponding elements are compressed in the temporal retinae relative to those in the nasal retinae [Shipley and Rawlings, 1970]. It should be reasonable to expect the horizontal misalignment of the eye’s optical elements to impact the asymmetry of the horizontal foveal correspondence. This impact of eye horizontally misaligned optical components on stereopsis and visual space geometry was confirmed in the geometric modeling in Turski [2018, 2020, 2023a] of the binocular system with the asymmetric eye (AE) model that comprised the horizontal displacement of the fovea from the posterior pole and the crystalline lens tilt relative to the eye’s optical axis. The results of those studies are briefly reviewed in the next section, which should help read the rest of the paper.

The study presented here provides the comprehensive geometric construction of spatial iso-disparity curves and the subjective vertical horopter in the binocular system complemented with the eye’s fovea and lens vertical misalignment in the AE model. The iso-disparity curves and the vertical horopter transformations are visualized in *GeoGebra’s* dynamic geometry simulations between different binocular postures. The transformed iso-disparity curves consist of families of ellipses and hyperbolas in which the zero-disparity horopter closely resembles the empirical horopter. Further, the perceived vertical horopter inclination from the true vertical agrees with experimental findings.

The results of this study provide new ideas for the physiologically motivated geometric conceptualization of binocular vision. In particular, it explains the functional role of vertical misalignment of the healthy human eye’s optical components.

### 2 2D Asymmetric Eye

The first asymmetric eye (AE) constructed in Turski [2018] comprised the horizontal displacement of the fovea from the posterior pole by angle *α* and the crystalline lens’ optical axis tilt relative to the eye’s optical axis by angle *β*, both measured at the nodal point, cf. Figure 1 (b). In the human population, angle *α* = 5.2*^◦^* is relatively stable, and angle *β* predominantly varies between *−*0.4*^◦^* and 4.7*^◦^*with the average value of *β* = 3.3*^◦^*.

**Figure 1:**
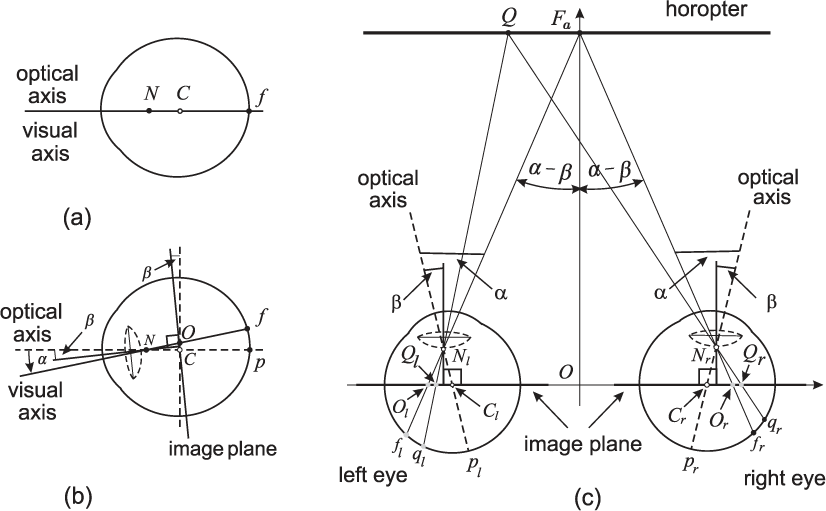
(a) The axially symmetric eye’s model of a single refractive surface schematic eye. (b) The right AE. *N* is the nodal point, *C* is the eye’s rotation center, *f* is the fovea, and *p* is the posterior pole. The angles *α* and *β* model horizontal misalignment of the human eye’s optical components. The image plane is parallel to the lens’s equatorial plane and passes through the eye’s rotation center. *O* is the projection of the fovea into the image plane—the optical center on the image plane. (c) The binocular system with the AEs fixates at *F_a_*, in which the coplanar equatorial lens planes are parallel to the coplanar image planes. This posture establishes the straight frontal horopter. The point *Q* on the horopter projects through the nodal points *N_r_* and *N_l_* to the corresponding retinal points *q_r_* and *q_l_* and to points *Q_r_* and *Q_l_* in the image plane. The fixation *F_a_* projects to the foveae *f_r_* and *f_l_* in the retina and the optical centers *O_r_* and *O_l_* in the image plane. Although *|f_l_q_l_|* ≠ *|f_r_q_r_|* on the retina, *|O_l_Q_l_|* = *|O_r_Q_r_|* on the image plane.

Figure 1 (a) presents the single refractive surface schematic eye, and Figure 1 (b) the AE model. In the clinical studies of eye asym-metry listed in the first section, different measuring methods (the Scheimpflug, Purkinje, and anterior segment optical coherence to-mography) were used to obtain the lens tilt relative to different reference axes (corneal topographic axis, line of sight, and papillary axis). Therefore, the values of the asymmetry angles defined geometrically in the AE have to be estimated from these clinical studies.

In the geometric theory developed in Turski [2020], the retinal correspondence of the binocular system with AEs is first constructed for the fixation corresponding to the abathic distance fixation char-acterized by the empirical horopter as a straight frontal line. In this posture, the eyes’ image planes and the lens’s equatorial planes are coplanar. It is shown in Figure 1 (c). This binocular posture is referred to as the eyes’ resting posture (ERP) because it numerically corresponds to the eyes’ resting vergence posture, in which the eye muscles’ natural tonus resting position serves as a zero-reference level for convergence effort [Ebenholtz, 2001]. The result is the symmet-ric distribution of the points in the image planes of the AEs that cover unknown asymmetric retinal correspondence under projection through the nodal points, explained in the caption of Figure 1.

For all other fixations in the horizontal plane, the horopters simulated in Turski [2020] in *GeoGebra’s* dynamic geometry environment consist of ellipses or hyperbolas resembling the empirical horopters. Importantly, the symmetric distribution on the image plane is preserved for all of these fixations, providing the first evidence that the asymmetry of retinal corresponding elements is caused by the hor-izontal misalignment of the eye’s optical components. The retinal correspondence asymmetry determines the shape of the longitudinal empirical horopter [Shipley and Rawlings, 1970, Nelson, 1977, Howard and Rogers, 2012].

Further, in Turski [2023a], the zero-disparity horopter is extended to the families of iso-disparity conic sections and applied to study in the framework of Riemannian geometry the global aspects of phenomenal spatial relations—visual space variable curvature and finite horizon. The global aspects of stereopsis are crucial in perception, as exemplified by the coarse disparity contribution to our impression of being immersed in the ambient environment despite receiving 2D projections of the light beams reflected by spatial objects [Barry, 2013].

The *GeoGebra* simulations in Turski [2023a] of iso-disparity lines in the ERP revealed that their distribution is invariant to changes of the AE parameters *α* and *β*. This invariance, demonstrated in Fig. 2 and Fig. 5 in that reference, shows that the distribution of iso-disparity lines is *universal*. It means that for any of the parameters *α* and *β* of the AE that uniquely prescribe the ERP (see Eq. (3) in Turski [2023a]), iso-disparity lines agree with this universal distribution. Moreover, the distance between iso-disparity lines can be scaled down to the cone photoreceptors separation in the foveal center or even down to hyperacuity for the optimal viewing distance of about 40 cm, see Fig. 6 in Turski [2023a].

**Figure 2:**
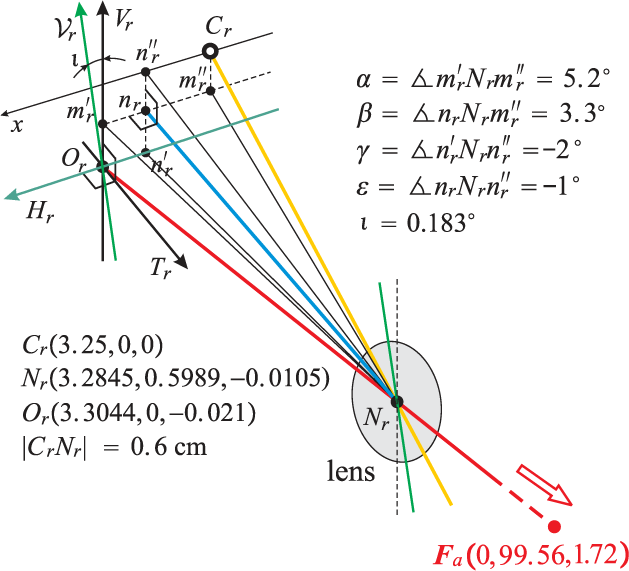
The right 3D AE in the ERP. The yellow line is the eye’s optical axis, the red line is the visual axis fixating *F_a_*, and the cyan line is the lens’s optical axis. The AE’s image plane coordinates consist of the axis *H_r_* of the intersection by the visual axis and the vertical axis *V_r_*, both attached at the optical center *O_r_*. They are complemented by the perpendicular coordinate *T_r_* parallel to the lens’s optical axis. These three axes in the ERP are parallel to the upright head coordinate axes. The details of other figures are discussed in the text.

The results obtained in Turski [2020, 2023a] firmly establish for the first time that the human eye misalignment of optical components results in the asymmetry of retinal corresponding elements. This asymmetry, in turn, impacts iso-disparity curve variation with the eye’s binocular fixation.

Although the spatial coordinates of retinal disparity (iso-disparity curves) provide the basic conceptualization of stereopsis, many different computational methods in its studies have been applied. One of the methods in Pollard et al. [1985] is based on salient features such as edges or corners, and another in Blake and Zisserman [1987] is based on higher-order properties of surfaces interpolated from data points to obtain the depth map. Still, many other computational techniques lead to the orientation, spatial frequency, or gradient disparities [Blakemore, 1970, Tyler and Sutter, 1979, Koenderink and van Doom, 1976, Rogers and Cagenello, 1989, Gåarding and Linde-berg, 1994]. Despite the contributions from these approaches to stereopsis, the recent claim in Lappin [2014] that retinal corresponding elements and disparity spatial coordinates (iso-disparity curves) are not necessary for stereoscopic vision is unconvincing as it is demonstrated by results obtained in my studies that were mentioned above.

### 3 3D Asymmetric Eye Model: The Criterion of Retinal Vertical Correspondence

Based on the clinical studies listed in the first section, the angles of *γ* = *−*2*^◦^*, *ε* = *−*1*^◦^* are assumed for the vertical asymmetry of the fovea and the lens, respectively, complementing *α* = 5.2*^◦^*and *β* = 3.3*^◦^* angles of horizontal asymmetry. For these asymmetry angles, the lens equatorial planes and, hence. the image planes of the AEs are coplanar for the fixation *F_a_*(0, 99.56, 1.72) expressed in centimeters in the upright head coordinate system—this is the ERP. Figure 2 shows the ERP of the binocular system perspective view of the right 3D AE near the optical center *O_r_* and the eye’s rotation center *C_r_* (3.25, 0, 0) in the image plane. The coordinates of *O_r_* and the nodal point *N_r_* are obtained in *GeoGebra’s* simulation for the values of the misalignment angles. The left AE is the mirror-symmetric in the median plane.

In Figure 2, the axis *H_r_* is the axis of retinal horizontal correspondence, i.e., the location of the corresponding points on the intersection of the coplanar image planes parallel to the lenses’ equatorial plane with the visual plane. Thus, the line *H_r_* is usually called the retinal horizon. The inclination of the visual plane passing through *O_r_*, *O_l_* and *F_a_* is *η* = arctan(1.741*/*99.56) = 1*^◦^*. The direction of retinal vertical correspondence *V_r_* differs from the vertical axis *V_r_* by the angle *ι* = 0.183*^◦^* constructed below for the AE’s misalignment angles. The coordinate axes *H_r_* and *V_r_* in the image plane are shown in green color.

Helmholtz [1867/1925] classic theory presumed that in the up-right head, binocular system fixating on a point in the intersection of the horizontal plane and the median plane, the meridians of the vertical retinal correspondence are not perpendicular to the retinal horizon. This lack of perpendicularity is known as *Helmholtz’s shear*. However, the main vertical meridians are not well defined in the human eyes because of the global eyeball asymmetry caused by the fovea displacement on the retina from the posterior pole by about 5*^◦^* measured at *N_r_*. This angle measured at the eye center is about 8*^◦^*. For this reason, in the geometric theory of the binocular system with AEs, the meridian criterion of verticality is replaced with the lens orientation criterion of verticality. Moreover, this is done for any fixation in the 3D binocular field, significantly extending Helmholtz’s theory.

Before I continue, I recall that in geometric theory in Turski [2020, 2023a], the eyes in the upright head are binocularly fixated on points in the horizontal plane, the image plane corresponding points are always located on the line intersecting the visual plane and the image plane, which is precisely the retinal horizon of the binocular system. In the study presented here, the eyes can fixate on the points of the 3D binocular field. Thus, in tertiary eyes postures, the ocular torsion twists the retina, resulting in nonzero torsional disparity. Although for a constant angular size of this disparity, its linear size is increasing with eccentricity, the binocular single vision is maintained in the peripheral region by sensory cyclofusion of up to 8*^◦^* supported by pe-ripheral large receptive fields [Guyton, 1987]. While one can measure fusion horopter [Amigo, 1974, Tyler, 1991, Harrold and Grove, 2015] to determine the horopter curve as its medial axis, it will not help my geometric modeling of stereopsis.

Here, the horizontal correspondence is again expressed on the in-tersection line of the visual plane and the image plane of the AE, initially in the ERP and then maintained in simulations for other fixations. This simple construction is allowed by the sensory cyclofusion range of 8*^◦^* in addition to the active compensation for the ocular torsion by the brain processing [Daddaoua et al., 2014] and the fact that the largest torsional disparity in the simulations is less than 1*^◦^*.

Because the main vertical meridian that serves as the basis for the verticality criterion is not well defined in the eye with the misaligned optical components, the vertical line in the equatorial plane of the lens substitutes the main meridian. This line, shown in Figure 2 in green that passes through *N_r_*, tilts when the lens undergoes both horizontal and vertical tilt and is always parallel to the image plane, defining the direction of vertical retinal correspondence.

The line of vertical retinal correspondence is constructed as follows. Let *M_r_* be the plane containing *n_r_*, *N_r_*, and the origin of the upright head coordinate system where *n_r_* is the orthogonal projection of *N_r_* into the image plane. The normal vector to *M_r_*, **v**, is parallel to the image plane and is perpendicular to the position vector **n***_r_*. Thus, **v** = (0.0105, 0, 3.2845). It defines the line of vertical foveal correspondence *V_r_* passing through *O_r_* (shown in Figure 2 as the green line) and tilted by *ι* = arctan(0.0105*/*3.2845) = 0.183*^◦^* relative to the objective vertical axes *V_r_* with its top in the temporal direction. Because of the mirror symmetry in the median plane in the ERP, the left AE’s vertical retinal correspondence is similarly formulated. Thus, the tilt between the lines of vertical retinal correspondence is 2*ι* = 0.366*^◦^*.

I end this section by presenting in Table 1 results obtained in *GeoGebra* for a couple of different vertical misalignment values of *γ* and *ε* for the constant values of *α* and *β* listed above. The distribution of iso-disparity curves was studied for three different values of *β* in Turski [2023a]. The table data include the corresponding fixation point *F_a_*, the tilt of the retinal apparent vertical *ι*, and the inclination of the visual plane *η*.

**Table 1:** Results for the different values of AE’s vertical angles *γ* and *ε* for *α* = 5.2*◦* and *β* = 3.3*◦*.

| $\gamma, \varepsilon$ | Fixation Point | $\iota, \eta$ |
| --- | --- | --- |
| $-1, -1$ | $F_a(0, 99.61, 0)$ | $0.183^\circ, 0^\circ$ |
| $-2, -1$ | $F_a(0, 99.56, 1.72)$ | $0.183^\circ, 1^\circ$ |
| $-2, -2$ | $F_a(0, 99.61, 0)$ | $0.366^\circ, 0^\circ$ |
| $-3, -1$ | $F_a(0, 99.55, 3.45)$ | $0.183^\circ, 2^\circ$ |
| $-3, -2$ | $F_a(0, 99.56, 1.72)$ | $0.366^\circ, 1^\circ$ |

The results obtained in this section and summarized in Table 1 should be noted for their simplicity. First, the retinal apparent vertical tilt increases by 0.183*^◦^* for each additional *−*1*^◦^*of the lens vertical tilt *ε*. Second, the inclination of the visual plane *η* is equal to *ε − γ*.

## 4 Construction of Vertical Horopter and Calculation of Ocular Torsion

### 4.1 Iso-disparity Conic Sections and the Vertical Horopter

Figure 3 explains the construction of the disparity conic sections and the vertical horopter for the fixation point *F* (*−*20, 159.56, 21.72) of the simulation in Figure 10, discussed below.

**Figure 3:**
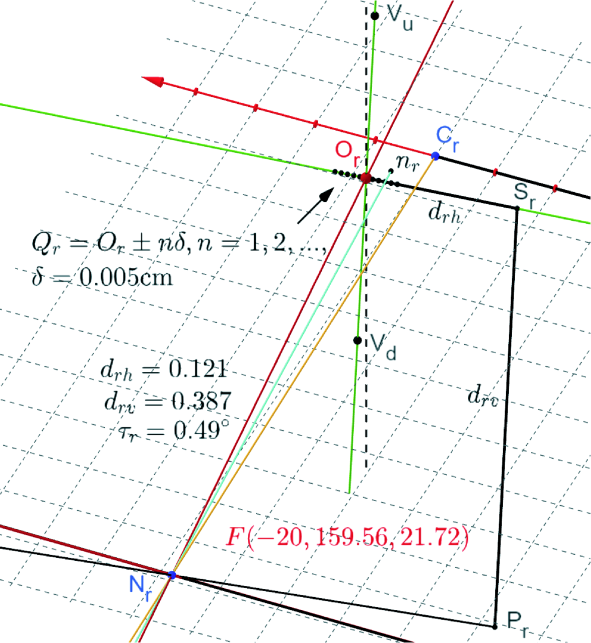
The construction of the iso-disparity hyperbolas and the vertical horopter for simulation in Figure 10 are shown for the right eye. The red line passing through the nodal point *N_r_* and the optical center *O_r_* is the visual axis, and the yellow line from *C_r_* and through *N_r_* is the optical axis. The cyan line through *N_r_* and *n_r_* is the lens’s optical axis perpendicular to the image plane. The fixation point is *F* (*−*20, 159.56, 21.72), and the horizontal and vertical coordinates relative to *O_r_* are listed. The green line through *O_r_* and *S_r_* is the foveal horizon with the horizontal disparity points placed 0.005 cm apart. The other green line through *O_r_* is the axis of vertical retinal correspondence—the axis *V_r_* in Figure 2—used to construct the vertical horopter, explained in the text.

Similarly, as in Figure 1 (c), the iso-disparity lines in physical space are first constructed for the ERP as follows. In Figure 3, the points *Q_r_* = *O_r_ ± nδ*, where *δ* = 0.005 cm and *n* = 1, 2,.., are shown on the foveal horizon line (green line through *O_r_* and *S_r_*). The *n*-th iso-disparity line is obtained for all pairs (*Q_r_, Q_l_*) such that *Q_r_ − Q_l_* = *nδ*. In particular, for *n* and *n* + 1, Figure 5 shows consecutive straight frontal is-disparity lines for the ERP of fixation point *F_a_*(0, 99.56, 1.72). Then, the hyperbolas in Figure 10 are obtained in the *GeoGebra’s* simulation after *F_a_* has changed to *F* (*−*20, 159.56, 21.72).

To construct the subjective vertical horopter, I choose two points *V_u_* and *V_d_* on the apparent vertical green line in the image plane passing through the optical center *O_r_* for the right AE in Figure 3. This line is the axis *V_r_* shown in Figure 2 indicating the foveal vertical criterion the 3D lens tilt prescribes. The points *V_u_* and *V_d_* are the same distance from the optical center. The same construction is carried out for the left AE. Two intersections of the projecting rays would define the vertical horopter, one for *V_u_* points and the other for *V_d_* points of the AEs. However, the intersection is not empty for fixations in the median plane, which is usually considered in experiments. Thus, it works for simulations in Figures 6, 7 and 11.

The vertical horopter is extended to the fixation’s central range of azimuthal angles by using the 3D property of this horopter, first emphasized by Amigo [1974], who studied the stereoscopic sensitivities of the retinal regions at the various elevations above and below the fovea. The vertical horopter’s extension for asymmetric fixations is approximated as follows. The two back-projected rays, one for each AE used for constructing the vertical horopter in the ERP, are replaced with cylinders. The other two rays intersect the cylinders, and the vertical horopter passes through the midpoints of the intersections. The cylinder’s radius determines the vertical horopter range of the azimuthal angles. In the simulations, I assumed a 1 cm radius at the abathic distance of about 100 cm and 30 cm above and below the fixation point.

### 4.2 Ocular Torsion and Torsional Disparity

Euler’s fundamental rotation theorem states that any two orientations of a rigid body with one of its points fixed differ by a rotation about an axis, specified by a unit vector, passing through the fixed point. The rotation matrix can be represented as *R*(*ϕ*, **n**), where the body rotates by angle *ϕ* about the direction given by the unit vector **n**. The *Euler-Rodrigues parameters ρ* = cos(*ϕ/*2), **e** = sin(*ϕ/*2)**n** produce the unit quaternion (cos(*ϕ/*2), sin(*ϕ/*2)**n**) and *Rodrigues’s vector*

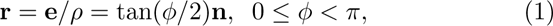

each representing the rotation by *ϕ* around **n**; see Piña [2011]. Note that **r***^−^*^1^ = *−***r** and (*−ϕ, −***n**) and (*ϕ*, **n**) represent the same vector. The ocular torsion simulation in the binocular system with the AE is briefly explained with the help of Figure 4.

**Figure 4:**
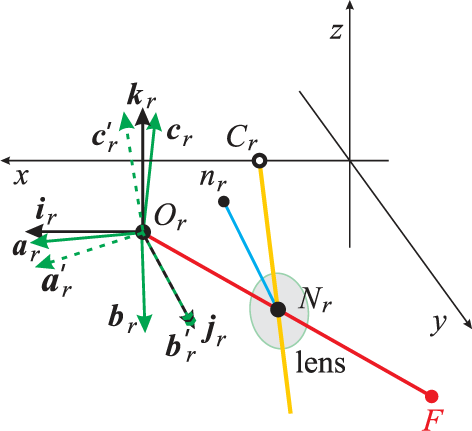
Schematic explanation of how the ocular torsion is defined and calculated for the right eye in the binocular system with the AE. The black frame agrees with the head frame at ERP. The green frame corresponds to the eye fixating at *F*. The green “primed” frame in dashed lines shows the rotated green frame in the plane containing the vectors **b***_r_* and **j***_r_* which overlays **b***_r_* with **j***_r_*. The geometric definition of the ocular torsion is ∠(**k***_r_*, 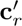).

In the ERP, the coordinate frames of the coplanar image planes of both eyes agree with the coordinate frame of the stationary upright head. In Figure 4, the black frame (**i***_r_*, **j***_r_*, **k***_r_*) represents the right AE of the ERP with its vectors along the respective axes (*H_r_, T_r_, V_r_*) shown in Figure 2. When the eyes change the ERP fixation to any other fixation *F*, the AE image planes are accordingly rotated. The new posture is then described by the green frame (**a***_r_*, **b***_r_*, **c***_r_*). The AE ocular torsion is calculated as follows. The green frame is rigidly rotated in the plane containing the vectors **b***_r_* and **j***_r_* such that **b***_r_* is overlaid with **j***_r_*. The resulting primed green frame (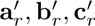) defines the ocular torsion by *τ_r_* = ∠(**k***_r_*, 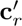). It establishes the geometrical definition of the eye ocular torsion for any binocular fixation executed from the ERP. The torsional disparity is the difference in ocular torsion between the right and the left eye.

Ocular torsion is usually considered the rolling motion of the gaze direction when the eye’s posture changes. Here, ocular torsion is geometrically defined as the rotation of the image plane’s perpendicular coordinate *T_r_* parallel to the lens’s optical axis. It is the best definition for the AE, which comprises the misalignment of the human eye’s optical components. It is justified by *|O_r_C_r_|* = 0.058 cm, cf. Figure 2.

The actual simulations of ocular torsion in the binocular system with AEs have been done in Turski [2023b], where the reader is referred for a comprehensive discussion and the impact on Listing’s law important in oculomotor perception. I note that the construction of the vertical horopter in that reference differs from the one presented here—the lens criterion of retinal vertical correspondence was not included in Turski [2023b]. However, the definition of ocular torsion for the eye’s misaligned optical components does not depend on the exclusion/inclusion of the lens-based criterion of the retinal vertical correspondence.

Here, I present for completeness one such simulation in Figure 5 that computes ocular torsion for the case simulated later in Figure 9. It explicitly describes Rodrigues’ vectors for Figure 9 simulation

**Figure 5:**
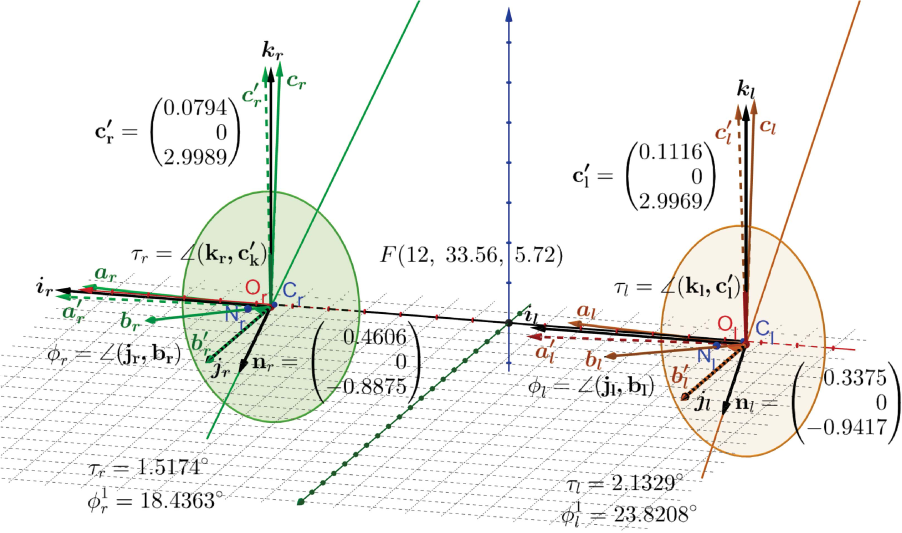
The black frames in the ERP (**i**_*r*_, **j**_*r*_, **k**_*r*_) and (**i**_*l*_, **j**_*l*_, **k**_*l*_) for the right and left AEs are attached at the image planes optical centers *O_r_* and *O_l_*. Their orientations agree with the upright head’s frame. The solid-colored frames are moving from the initial position of black frames for a given fixation *F*. The dashed-colored frames are obtained by rotating the solid-colored frames by moving **b**s vectors onto the **j**s vectors. The rotation axes are given by the vector **n**_*r*_ and **n**_*l*_, with the angle of rotations *ϕ_r_* and *ϕ_l_*, are contained in the co-planar image planes of the ERP, parallel to the head frontal plane. The ocular torsion of each AE is *τ_r_* = 1.52 for the right eye and *τ_l_* = 2.13 for the left eye.

**Figure 6:**
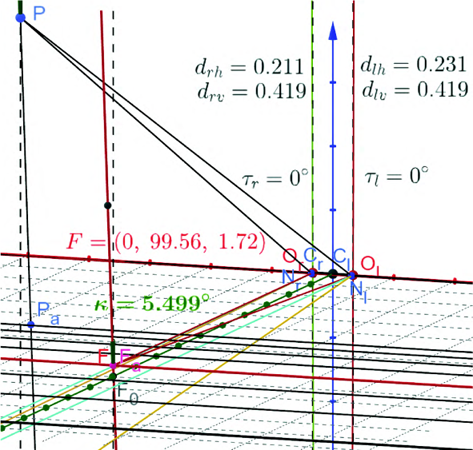
The ERP with the fixation at *F_a_* showing the iso-disparity frontal lines in the visual plane and the vertical horopter tilted by *κ* = 5.5*^◦^* relative to the objective vertical. The longitudinal and vertical horopters passing through *F_a_* are shown in red. The point *F*_0_ is the vertical projection of *F_a_* into the head’s transverse plane containing the eyes’ rotation centers (marked with the grid). The details of other figures are discussed in the text.

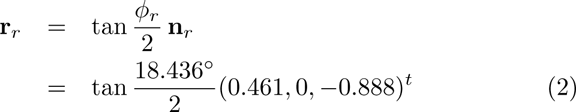

for the right eye and

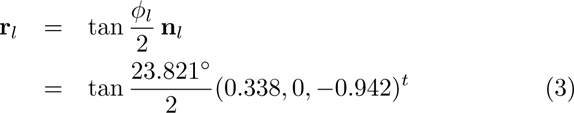

for the left eye. Here, ‘*t*’ denotes the transpose. These Rodrigue’s vectors rotate the corresponding frame vectors discussed in Figure 5, resulting in the ocular torsion for each eye.

## 5 Simulations of Retinal Correspondence Spatial Coordinates

In this section, I visualize the longitudinal disparity spatial coordi-nates (iso-disparity curves) and the subjective vertical horopter in the binocular system with 3D AE using the simulation in *GeoGe-bra’s* dynamic geometry environment. The results also include the horizontal and vertical disparities and the ocular torsion for each binocular fixation of the AEs.

The iso-disparity curves and the subjective vertical horopter are precisely defined in the binocular system with AEs only for fixations in the median plane. The extension of the iso-disparity curves, especially to tertiary fixations, was discussed in Section 3 and constructed in Section 4.1. Similarly, the vertical horopter should be extended by accounting for Panum’s fusional region of the empirical vertical horopter measured in Amigo [1974]. A simple extension of the vertical horopter as a straight line used in this section’s simulations was also constructed in Section 4.1. Moreover, the computations of ocular torsion were explained in Section 4.2.

### 5.1 Disparity Spatial Coordinates for ERP

The fixation point *F_a_* of the ERP of the stationary upright head is uniquely determined by the assumed parameters of the binocular system with the AE. Then, *F_a_*’s coordinates in centimeters are (0, 99.56, 1.72), corresponding to the average abathic distance of empirical horopters of about 1 meter. The simulated iso-disparity conic sections in the visual plane are straight frontal lines in the ERP. Also, in this posture, the vertical horopter is a straight line tilted from the true vertical by *κ* = 5.5*^◦^*with its top away from the head, shown in Figure 6.

This figure shows the disparity lines for the proximal relative disparity of *δ* = 0.005 cm, explained in Figure 3. The point *P*, shown in Figure 6, is projected into the AE image planes through the nodal points. The right image plane projection is (*d_rh_*(*P*), *d_rv_*(*P*)) and the left image plane projection is (*d_lh_*(*P*), *d_lv_*(*P*)) in the coordinates *H_r_* and *V_r_* and *H_l_* and *V_l_*, respectively. From the simulation computed in *GeoGebra* and displayed in Figure 6, the horizontal disparity of *P* is

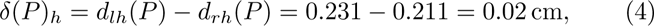

but the vertical coordinates *d_rv_* and *d_lv_* are the same, so *δ*(*P*)*_v_* = 0.

The projection of *P* into the visual plane along the line parallel to the vertical horopter, *P_a_*, is exactly on the fourth consecutive disparity line down from the horopter (red frontal line through *F_a_*). Because the horizontal retinal relative disparity value between each consecutive disparity line is 0.005 cm, the spatial disparity of *P* agrees with the retinal horizontal disparity *δ*(*P*) *_h_* = 4(0.005) = 0.02 cm. The horizontal disparity for crossed disparities is always considered positive, and for uncrossed disparities, it is negative. The ocular torsion, *τ_r_*, and *τ_l_*, computed in this simulation, is zero, such that the torsional disparity *δ_t_* vanishes in this posture.

### 5.2 Disparity Spatial Coordinates for Eyes Secondary Postures

The iso-disparity curves and the vertical horopter are shown in Figure 7 for the fixation point *F* that is shifted from ERP by the down-shift from *F_a_*(0, 99.56, 1.72) to *F* (0, 99.56*, −*21.72).

**Figure 7:**
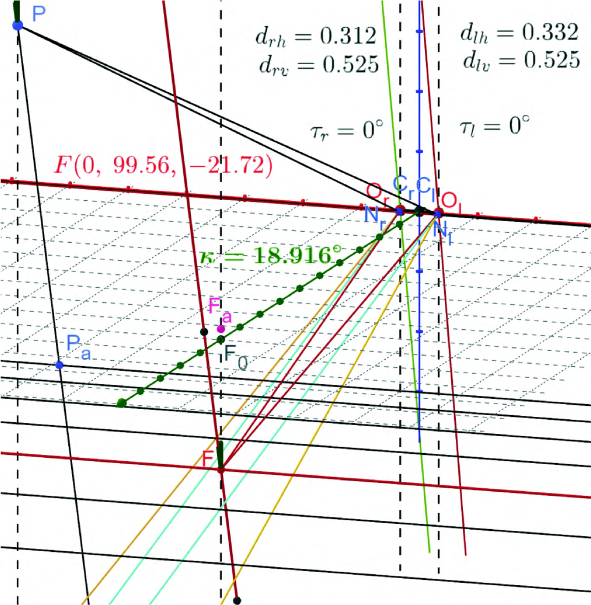
Under the true vertical shift of the fixation point from the ERP, the iso-disparity lines move with the visual plane and remain straight frontal lines. The subjective vertical horopter is tilted from the true vertical direction by *κ* = 18.9*◦*. See the text for a detailed discussion.

This figure shows that the iso-disparity lines move with the visual plane remaining straight and frontal. Further, the subjective vertical horopter is tilted in the midsagittal plane top-away by *κ* = 18.9*^◦^* at the fixation point from the objective vertical line shown by a dashed line through *F_a_*.

Under this down-shift movement of *F_a_*, the horizontal disparity of point *P* shown in this figure (this point is different than point *P* in Figure 6) changes to

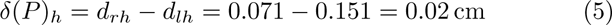

but its vertical coordinates *d_rv_* and *d_lv_* are the same, so *δ*(*P*) *_v_* = 0. Further, the ocular torsion vanishes for each eye, so the torsional disparity remains zero after the vertical shift of *F_a_*.

The projection ray of *P* onto *P_a_* in the visual plane is also parallel to the subjective vertical horopter in this posture. Again, in Figure 7, the projection of *P* along the subjective vertical direction into the visual plane, *P_a_*, is on the disparity line of disparity value 0.02 cm. It agrees with the horizontal disparity value obtained from the differences of the projections in the coordinates in the right and left eye image planes and shown in Eq. (5). It demonstrates that under the true vertical shift of the fixation point from the ERP, the visual space tilts together with the tilt of the subjective horopter by the same amount. The next secondary posture is produced by shifting *F_a_*(0, 99.56, 1.72) horizontally to *F* (40, 99.56, 1.72), shown in Figure 8.

**Figure 8:**
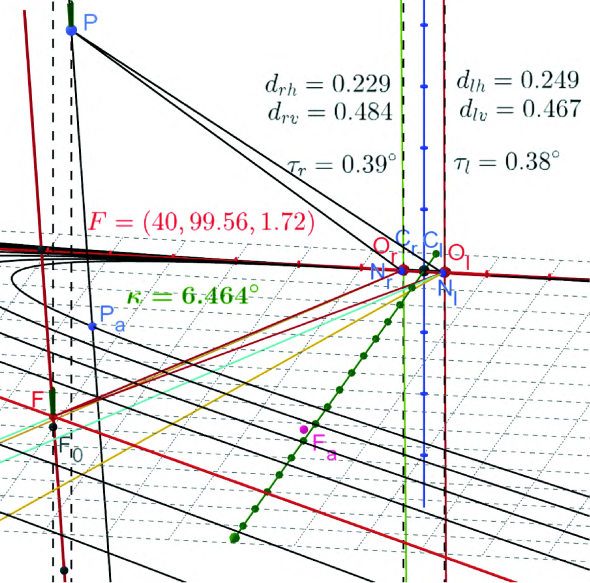
Under the horizontal shift of the fixation point from the ERP, the disparity lines in the visual plane transform into hyperbolas. The subjective vertical horopter is tilted from the true vertical direction by about *κ* = 6.5*^◦^*.

Now, the iso-disparity lines in the image plane change into hyperbolas for this secondary posture. Further, the subjective vertical horopter is tilted top-away by about *κ* = 6.5*^◦^* in the plane *−x* + 0.5*y* = 6.8 containing the subjective vertical horopter and the objective vertical line (dashed line). Using the same argumentation as before, we see that the line through *P* and *P_a_*, where *P* is chosen such that the longitudinal spatial disparity value of 0.02) cm agrees with the retinal horizontal disparity value of *δ*(*P*) *_h_* = 0.02 cm. In contrast to the vertical secondary posture, the vertical and torsional disparities are small but not zero after the horizontal shift: *δ*(*P*) *_v_* = 0.017 cm and *δ_t_* = *τ_r_ − τ_l_* = 0.01*^◦^*.

### 5.3 Disparity Spatial Coordinates for Eyes Tertiary Postures

Figures 9 and 10 show the disparity ellipses and hyperbolas for the tertiary postures when the resting eyes fixation point *F_a_* is shifted to positions by changing all three components of the fixation point.

**Figure 9:**
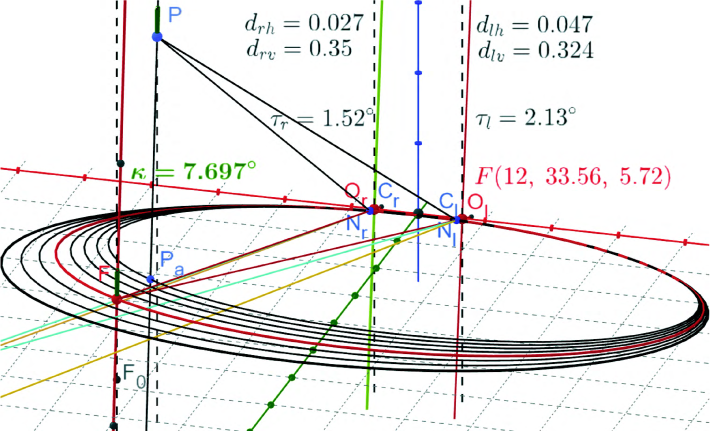
The iso-disparity curves are ellipses, and the vertical horopter is tilted *κ* = 7.7*^◦^* for the fixation of the distance less than the abathic distance.

In Figure 9, for the fixation at *F* (12, 33.56, 5.72), the horizontal disparity value is *δ*(*P*) *_h_* = 0.02 cm, the vertical disparity value is *δ*(*P*) *_v_* = 0.026 cm and the torsional disparity value is *δ_t_* = *τ_r_ − τ_l_* = *−*0.61*^◦^*, cf. Figure 5. Also, the subjective vertical horopter is tilted in the plane 1.3*x* + 0.12*y* = 20 by *κ* = 7.7*^◦^* top-away relative to the true vertical.

**Figure 10:**
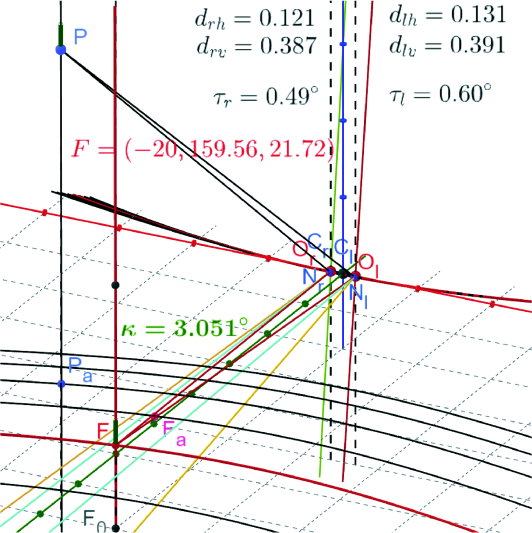
The disparity curves are hyperbolas, and the vertical horopter is tilted *κ* = 3.05*^◦^* for the fixation of the distance greater than the abathic distance. Note that the hyperbolic branches that pass through the nodal points *N_r_* and *N_l_* are not part of disparity curves.

In Figure 10, for the fixation at *F* (*−*20, 159.56, 21.72), the horizontal disparity value is *δ*(*P*) *_h_* = 0.01 cm, the vertical disparity value is *δ*(*P*) *_v_* = *−*0.004 cm and the torsional disparity value, *δ_t_* = *−*0.11*^◦^*. Finally, the subjective vertical horopter is tilted bottom-away in the plane 4.8*x* + 2.3*y* = 265 by *κ* = 3.05*^◦^*.

## 6 Discussion of the Simulated Eyes Postures

The first conclusion from the simulations of the eyes’ postures is that the equality of the retinal disparity obtained from the projections onto the image planes and the value obtained from the spatial iso-disparity coordinates shows that the perceived vertical agrees with the subjective vertical horopter. This conclusion disagrees with the results in Cogan [1979].

The simulation in Figure 6 shows that the spatial disparity in the ERP is organized by true vertical planes passing through the frontal disparity lines. For any point in the binocular field, the vertical disparity is zero, and the horizontal disparity value is given by the frontal disparity line above which the point is located on the vertical plane. The backward tilt of the vertical horopter measured in Amigo [1974] for two subjects were about 6*^◦^* and 8.5*^◦^* for the observation distance of 1 m and 12*^◦^* and 16.5*^◦^* for the observation distance of 3 m. Figures 6 and the simulation in the next Figure 11 show a reasonable agreement with Amigo’s results, given that he had not specified the misalignment of optical components of the subject’s eyes. These results also agree with the experimental results in Siderov et al. [1999]. Thus, my results do not agree with Helmholtz and others [Nakayama, 1977, Cogan, 1979] regarding the magnitude of the vertical horopter tilt.

**Figure 11:**
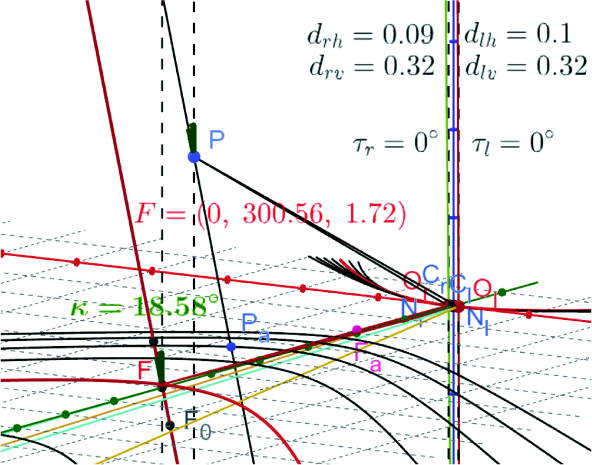
The iso-disparity curves and the vertical horopter simulated for the fixation *F* (0, 300.56, 1.72)

For the true vertical shift of the fixation point from the ERP shown in Figure 7, the objective binocular space (and vertical planes above disparity lines) tilts by the same amount as the vertical horopter tilts at the fixation point. This visual space is seen subjectively as the same before the fixation point was shifted. In particular, the vertical and torsional disparities remain zero.

These results indicate that the curvature of visual space vanishes for those eyes’ postures, extending the 2D results in Turski [2023a] for the horizontal visual plane to the 3D results for these two postures. Moreover, they clearly show that the vertical disparity is not a binocular cue, agreeing with the general evidence from the random-dot-stereograms Julesz [1971] and Wheatstone stereoscope Wheatstone [1838].

The situation is more complicated for the simulating horizontal shifts from the ERP shown in Figure 8 for the secondary position with the eyes fixating on *F* (40, 99.56, 1.72). Although the values of the vertical and torsional disparities are small and the straight line at the fixation well approximates the hyperbola, it can affect our impression of being immersed in the ambient environment.

For the eyes shifted vertically or horizontally when both are in secondary positions, their disparity curves are markedly different after each shift, as shown in Figures 7 and 8. This should be expected because the stereopsis is mainly related to the eyes’ lateral offset. My observation is that I use up-and-down eye movement more often without the head movement than when I move my eyes to the left or right. Is this behavioral dissimilarity connected to preserving the straight frontal disparity lines after vertical shifts and transforming them into hyperbolas after horizontal shifts? This question is worth investigating.

The one common feature of the vertical horopter orientations in Figures 8 – 10 is that under the change in fixation point, the vertical horopter’s tilt is specified in the computed vertical plane in each simulation. This aspect was never considered in experimental observations.

In the tertiary posture simulations shown in Figures 9 and 10, the geometry of visual space is much more complicated, even under the assumption of a straight-line vertical horopter. It should be noted that in Cooper et al. [2011], the numerical analysis produced convex vertical horopters. The authors concluded that the vertical horopter is adaptive for perceiving convex, slanted surfaces at short distances. However, the numerical methodology employed to discuss the vertical horopter curvature uses the Vieth-Müller circle (VMC) and the Hering-Hillebrand deviation coefficient *H* introduced in Ogle [1932]. However, each VMC and Ogles’ models of empirical horopter pass through the eye’s rotation centers rather than the nodal points; see Turski [2016, 2018] and for a more comprehensive critical discussion, see Turski [2023c].

One way to resolve the issue of the vertical horopter shape could be an experimental study similar to Amigo [1974] or Siderov et al. [1999] but with tested eye’ misalignment of the optical components and supported by the numerical methodology based on the AE model. Nevertheless, the results add to other contentions about the relevance of Luneburg’s assertion of the constant hyperbolic curvature visual space [Luneburg, 1947, Blank, 1957].

## 7 Conclusions

The iso-disparity lines and the vertical horopter provide our three-dimensional visual experience with the organizing geometric principles underlined by the anatomy and physiology of the binocular system. This assertion is emphasized by the author’s studies of the geometric constructions supported by computer visualizations in the binocular system comprising the eye model that incorporates the healthy human eye’s misalignment of the optical components. The asymmetric eye (AE) defined in Turski [2018] and extended in the later studies consists of the fovea not located on and the lens tilted away from the eye’s optical axis.

The study presented here extends their previously considered horizontal misalignment with the vertical components. The consequences and functional roles of the horizontal misalignment of the eye’s optical elements for the quality of vision are well known—the fovea’s anatomical displacement from the posterior pole and the asphericity of the cornea contribute to optical aberrations and the lens tilt partially compensates for these aberrations.

The neglected aspect of horizontal misalignment of the eye’s optical components involves its impact on the horizontal foveal correspondence, which organizes visual perception of objects’ form and location in 3D space, i.e., stereopsis and spatial relations. Recall that for a small retinal area in one eye, there is a corresponding unique area in the other eye such that both share one subjective visual direction. The results obtained in Turski [2023a] and reviewed here firmly establish for the first time that the universal distribution of the iso-disparity lines to the eye’s misaligned optical components results in the asymmetry of the retinal corresponding elements. This asymmetry, in turn, impacts the iso-disparity curve’s shape variation with the eye’s binocular fixation.

The transformations of iso-disparity curves between different eyes’ postures, visualized in GeoGebra’s dynamic geometry simulations, consisted of families of ellipses or hyperbolas depending on the eye’s horizontal misalignment. In particular, the horopter’s shapes resemble the shapes of empirical horopters during transformations. Further, ocular torsion for eyes’ postures was computed using Euler’s rotation theorem in Rodrigues’ (rotation) vector framework by closely approximating the eye’s rolling motion about the gaze direction. For the comprehensive study of ocular torsional motions during binocular fixations, the reader is referred to Turski [2023b].

This study’s original contributions include an examination of the consequences and functional roles of vertical misalignment of the eye’s optical components. It is explained in the following findings: The classic Helmholtz theory, which states that the subjective vertical retinal meridian inclination to the retinal horizon explains the backward tilt of the perceived vertical horopter, loses its relevance when the eye’s optical components are misaligned. Instead, the vertical lens’ tilt provides the retinal vertical criterion that explains the ex-perimentally measured vertical horopter’s inclination.

